# Intubation and mechanical ventilation lead to an extensive expansion of VAP-associated bacterial species in respiratory secretions of ventilated pigs

**DOI:** 10.64898/2026.10.14.759075

**Authors:** Lara Spring, Dorina Haider, Andrea Hoffmann, Fabian Zink, Martin Hessling, Peter Radermacher, Franziska Münz, Ohad Sharon, Magnus Scheer, Maximilian Feth, Thomas Kapapa, Barbara Spellerberg

## Abstract

Ventilator associated pneumonia (VAP) remains one of the most critical nosocomial infections in intensive care medicine. Between 20-50% of ventilated patients develop VAP during treatment, often leading to long term sequelae and death, despite successful therapy of their underlying disease. VAP is typically caused by Gram-negative bacteria that can previously be detected on the mucosal membranes of the oral and upper respiratory tract. To assess and quantify bacterial colonization of the respiratory tract following intubation and ventilation a well-established pig model was used.

**Methods:** After induction of anaesthesia and endotracheal intubation pigs underwent mechanical ventilation continuously for up to three days. To assess bacterial colonization of the upper respiratory tract, samples of respiratory secretions were taken at day 0, 1 and 2 directly above the cuff outside of the endotracheal tube (ETT) and by aspiration of tracheal secretions through the ETT. Quantification of bacteria was carried out by serial dilutions and CFU/ml determinations on sheep blood agar and MacConkey agar plates. Species identification was achieved by 16S-RNA gene sequencing.

**Results:** Respiratory samples of 19 pigs were analysed and showed a sharp increase of predominantly Gram-negative organisms in the respiratory material sampled above the cuff from day 0 to 1. Starting at a median of 7.3 × 10^3^ immediately after intubation numbers increased to a median of 2.0 × 10^7^ Gram-negative bacteria at day 1 and 3.2 × 10^7^ at day 2. Gram-negative species that were detected included *Pseudomonas aeruginosa, Acinetobacter baumanii, Enterobacter aerogenes, Klebsiella pneumoniae* and other *Enterobacteriaceae*. Tracheal secretions contained much less bacteria. While no Gram-negative bacteria could be cultured at the beginning of the experiment in 18 of 19 animals, colonization with Gram-negative bacteria gradually increased and was found in 7 of 19 and 9 of 14 animals on day 1 and day 2 respectively. Interestingly, in the vast majority of colonized tracheal secretions the identical Gram-negative species had previously been detected in secretions collected > 24h earlier at the cuff.

**Conclusion:** A massive increase of the concentration of Gram-negative bacteria occurred at 24h of ventilation in mucosal secretions above the cuff. Detected species were the same Gram-negative bacteria that are frequently responsible for human infections, suggesting that especially the accumulation of bacteria in secretions above the cuff may be crucial in the development of VAP.

## Introduction

Ventilator associated pneumonia (VAP) is a major reason for life threatening nosocomial infections in intensive care medicine. It is defined as a lower respiratory tract infection occurring in mechanically ventilated patients after ≥48 hours of endotracheal intubation (Torres *et al*., 2017) (Howroyd *et al*., 2024). The placement of an endotracheal tube (ETT) is the single most important risk factor for the development of VAP (Hunter, 2012). Endotracheal intubation circumvents natural airway defences by eliminating mucociliary clearance mechanisms and by creating a continuously open access to the lower respiratory tract. Depending on additional risk factors such as immunosuppression, preexisting pulmonary disease, trauma and in particular traumatic brain injury (Hu *et al*., 2017), between 20-50% of ventilated patients develop VAP during ICU treatment, often leading to long term sequelae and death (Lynch, 2001) (Wu *et al*., 2019). The estimated risk for the development of VAP lies between 2 to 16 episodes per 1000 ventilator days (Rosenthal *et al*., 2012) and increases with the duration of ventilation. Although the assessment of mortality solely attributable to VAP is difficult in multimorbid patients, several reports estimate it to lie between 3 % and 25.6% (Bekaert *et al*., 2011) (Melsen *et al*., 2011). Even in patients surviving VAP a considerable proportion suffer from long term sequelae such as impaired pulmonary function. VAP increases hospital stays by four to nine days, representing a major factor in health care costs, as it is one of the leading nosocomial infections (Kollef *et al*., 2012). Most often VAP is caused by Gram-negative bacteria (Restrepo *et al*., 2013) that can previously be detected on the mucosal membranes of the oral and upper respiratory tract and the microbiome of the lungs. Especially the group of ESKAPE pathogens (*Enterococcus faecium, Staphylococcus aureus, Klebsiella pneumoniae, Acinetobacter baumannii, Pseudomonas aeruginosa* and *Enterobacter* species), which is causing 80% of VAP, present with high antibiotic resistance rates (Low *et al*., 2026) (Giantsou *et al*., 2005). VAP accounts for half of the prescribed antibiotics in ICU settings, adding immensely to antibiotic pressure. Since mortality is significantly increased in infections involving resistant bacteria, high resistance rates are a major threat to favorable clinical outcome and survival of patients. Overall, the ESKAPE pathogens rank among the bacterial species most frequently associated with fatal infectious outcomes, underscoring their clinical importance (Antimicrobial Resistance, 2022).

A crucial factor in the development of VAP is the interference of endotracheal intubation with natural airway defense mechanisms. These include the elimination of nasal filtration and humidification of air during inspiration and a suppressed cough reflex. Mucociliary clearance is severely impaired (Sackner *et al*., 1975) (Konrad *et al*., 1994) and subglottic pooling of respiratory secretions and biofilm formation in the endotracheal tube provide an ideal environment for aberrant bacterial growth (Gil-Perotin *et al*., 2012). Necessary intensive care interventions like sedation, positive pressure ventilation and the positioning of the patient during ventilation further exacerbate this situation (Lynch, 2001). The impaired natural clearance facilitates a microbial shift in the oral flora, which helps bacterial pathogens that are not typical for the oral microbiota, in particular Gram-negative bacteria from the gastrointestinal tract, to become colonizers of the oral mucosal surfaces before gradually infiltrating the respiratory tract. These specific alterations of the respiratory microbiome have clearly been demonstrated by bacterial cultures as well as 16S-RNA based diagnostic methods (Ewig *et al*., 1999) (Kelly *et al*., 2016). Importantly, key features of respiratory colonization, such has bacterial burden and enrichment of Gram-negative gut associated bacteria have been shown to predict outcomes in critically ill patients (Dickson *et al*., 2020).

To quantify how intubation and ventilation alter the residential flora of the upper respiratory tract we determined the bacterial load in respiratory secretions collected at the cuff and in the trachea immediately at intubation and following 24 and 48 hours of ventilation in an experimental pig model. Since patients with traumatic brain injury (TBI) carry a particular high risk for the development of VAP (Hu *et al*., 2017), a TBI model was selected for investigating alterations of the microbial flora.

## Methods

### Large animal model

The study was conducted in a long-term ventilation pig model that was carried out at the Institute for Anesthesiological Pathophysiology and Process Engineering of the University Clinic Ulm. The current experiment is part of a greater experimental series on anesthetized, mechanically ventilated and (neuro)surgically instrumented pigs, which undergo combined acute subdural hematoma (ASDH) and hemorrhage and subsequent “TBI-targeted” intensive care with a total duration of mechanical ventilation of approx. 54-60 hours, thus enabling the investigation of bacterial species in respiratory secretions over at least two days. Moreover, using this experiment allowed for avoiding additional experimental animals according to 3R animal experiment principles (3R=Replacement, Reduction, and Refinement). All experiments were approved by the Animal Care Committee of the University of Ulm and the Federal Authorities for Animal Research (Regierungspräsidium Tübingen; Reg.-Nr. 1559, approval 29 October 2021) and were conducted following standards of the National Institute of Health Guidelines and the European Union Directive 2010/63/EU. The respiratory ventilation experiments were performed on 19 German large white pigs of either sex with ages ranging from 22 and 24 weeks and an average weight of 82 kg. 5 of 19 animals had to be euthanized prior to 48h of ventilation due to circulatory failure occurring during the main experiment.

### Collection of respiratory samples

After induction of anesthesia with propofol and ketamine and subsequent endotracheal intubation, the pigs underwent continuous volume-controlled mechanical ventilation under total intravenous anesthesia (TIVA) for up to 60 hours as detailed in (Abele *et al*., 2026). Samples of tracheal secretion were obtained through introduction of a suction catheter through the ETT and placement of the catheter a few cm just below the ETT. Pharyngeal secretions from the cuff area were collected directly above the cuff by placing the catheter outside of the ETT for aspiration. Afterwards the secretions were dissolved in 50ml of a sterile 0.9% sodium chloride solution. Collection of aspiration samples was performed immediately after intubation as well as after 24 hours and 48 hours of ventilation.

### Quantitative sample analysis

Tracheal and cuff secretion samples were serially diluted in sterile phosphate-buffered saline (PBS) up to a dilution of 10^−6^ and plated on sheep blood agar as a non-selective universal medium and on MacConkey agar as a selective agar for Gram-negative bacteria. Bacteria were incubated for 24 h at 37°C (with 5% CO_2_ for sheep blood agar and under aerobic conditions for MacConkey agar). After incubation, colony-forming units (CFU) were counted, and bacterial concentrations were calculated as CFU/ml based on corresponding dilution factor for both types of secretion samples obtained at 0, 24 + 1, and 48 + 1 h after intubation.

### Qualitative sample analysis

To identify the different bacterial species in the collected secretions, colonies retrieved during the previous quantitative analysis were subcultured on sheep blood agar to obtain pure cultures. Genomic DNA was isolated from the pure cultures using GenoLyse VER 1.0 Extraction kit (Bruker, Nehren, Germany) according to the manufacturer’s instructions. The bacterial 16S rRNA gene was amplified by PCR using primers 27F (5’-3’: AGAGTTTGATCCTGGCTCAG) and 1482R (5’-3’: ACGGGCGGTGTGTACAAG), which target conserved regions flanking variable sequences within the approximately 1,500 bp-long gene. The 16S rRNA gene is present in all bacteria and encodes for the rRNA of the 16S-unit of the bacterial ribosome. It contains both highly conserved regions, enabling primer binding, as well as hypervariable regions that allow taxonomic identification at the genus and species level. PCR amplification was performed using a *Taq* polymerase (Roche) under standard conditions. Briefly, an initial denaturation at 95°C for 5 min was followed by 30 amplification cycles of 1 min at 95°C, 30 s at 50°C, 90 s at 72°C, and a final elongation of 10 min at 72°C. The resulting amplicons were purified using the NucleoSpin® Gel and PCR Clean-up kit (Macherey-Nagel, Düren, Germany) according to the manufacturer's instructions. Purified PCR products were sent to Microsynth SeqLab Laboratories (Göttingen, Germany) for Sanger sequencing. Obtained sequences were analyzed using the Basic Local Alignment Search Tool (BLAST) at the NCBI server (hops://blast.ncbi.nlm.nih.gov/Blast.cgi). Sequence identities of at least 95% were considered indicative for same species.

### Statistical analysis

Statistical analysis was performed using GraphPad Prism 9.5.0. Due to the high interindividual variability and the non-normal distribution of data according to the performed Shapiro-Wilk test (p < 0.05), a nonparametric method was chosen. To determine general quantitative differences between measuring points the Friedman rank sum test was conducted. Since the inclusion of dropouts is not possible in this test, only 14 animals were factored in for the inferential statistical analysis, whereas the descriptive analysis still contains all 19 pigs. After receiving positive outcomes regarding a level of significance of 5%, further post-hoc analysis was performed using the Wilcoxon Matched-Pairs Signed-Rank Test to compare specific timepoints. Subsequently, p-values were adjusted to minimize alpha-error by applying a Holm-Bonferroni-correction. Results with a p-value of less than 0.05 were considered statistically significant.

## Results

### Quantitative analysis of bacterial species in respiratory secretions

To assess the effects of prolonged intubation and ventilation on the bacterial load found in the respiratory tract, a quantitative analysis of bacteria recovered from respiratory secretions collected at the cuff and in the lower trachea was performed. During the 48 hours of ventilation a highly significant rise in the total number of bacteria and in particular a rise of Gram-negative bacterial species was observed at the cuff (Fig.1).

**Figure 1:**
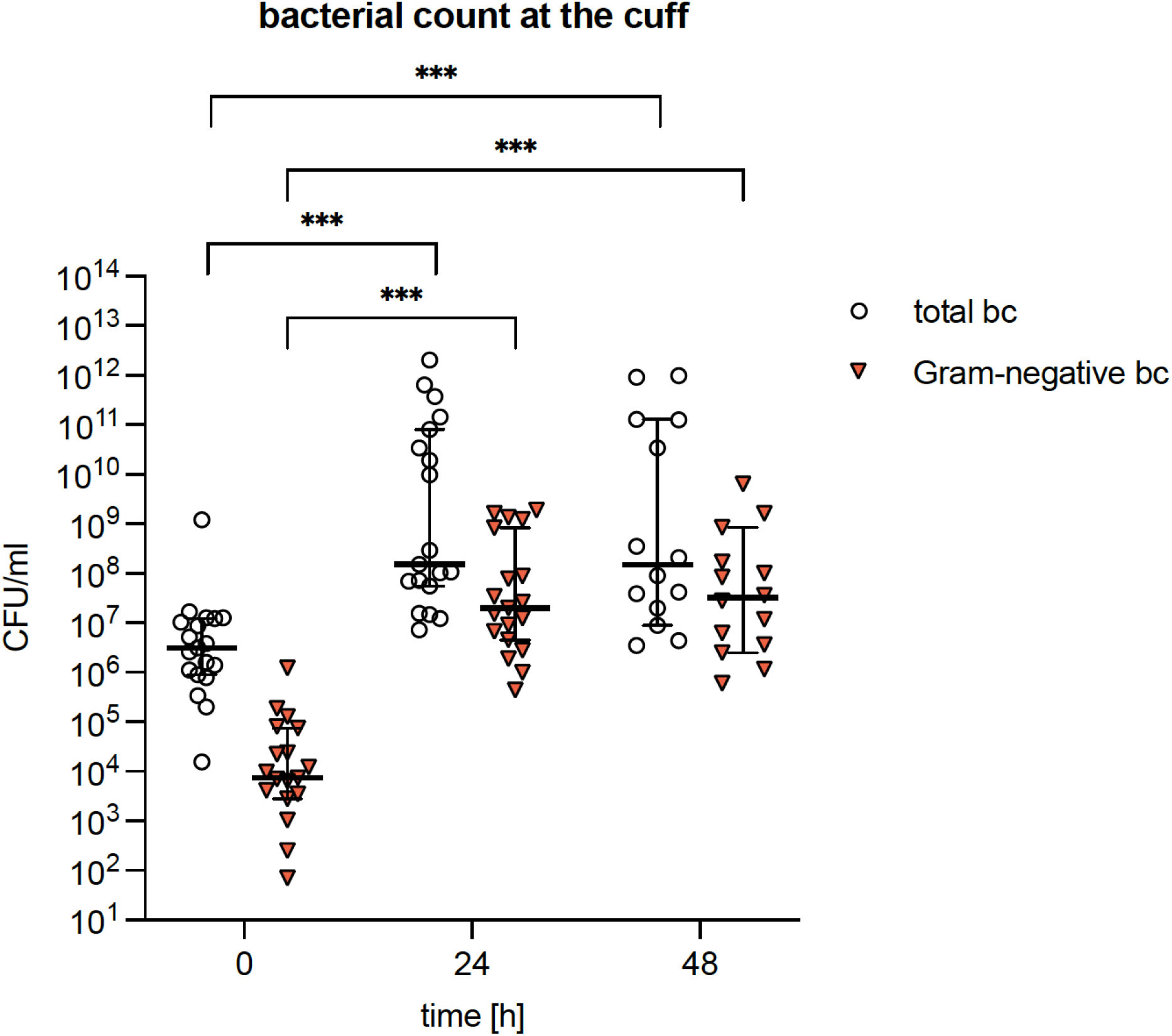
Depicted is the quantitative assessment of total bacterial counts (bc) and Gram-negative organisms collected in respiratory secretions at the cuff. Samples were taken immediately after intubation and after 24 + 1 hours (day1) and 48 + 1 hours (day2) of mechanical ventilation. CFU/ml determinations were carried out through diluting and cultivating respiratory samples on blood agar plates for total bacterial counts and MacConkey agar plates for growth of Gram-negative bacterial species. Shown are median bacterial counts and significant differences (p < 0.05) as determined by Wilcoxon Matched-Pairs Signed-Rank Test.

At the time of intubation total bacterial counts were 3.1 × 10^6^ CFU/ml multiplying to 1.5 × 10^8^ CFU/ml within 24 hours. As for Gram-negative species, the bacterial count was 7.3 × 10^3^ CFU/ml at the start of the experiment and reached 2.0 × 10^7^ CFU/ml at 24h and 3.2 × 10^7^ at 48h, which represents an increase of more than 3 orders of magnitude within one day. Unsurprisingly, these results, meaning the difference between 0 and 24 hours as well as 0 and 48 hours in both groups, were highly significant in Wilcoxon rank-sum tests. A similar development was observed in tracheal secretions. At the time of intubation, the median of total bacterial counts was very low at 10 CFU/ml for the total bacterial count and 0 CFU/ml for Gram-negative species. No significant difference in bacterial counts was noted for the first 24h, but after that timepoint total bacterial counts and Gram-negative bacterial counts rose significantly (Fig. 2). At 48 hours the total amount of bacteria was 7.6 × 10^3^ CFU/ml and the Gram-negative bacterial count reached 2.3 × 10^2^.

**Figure 2:**
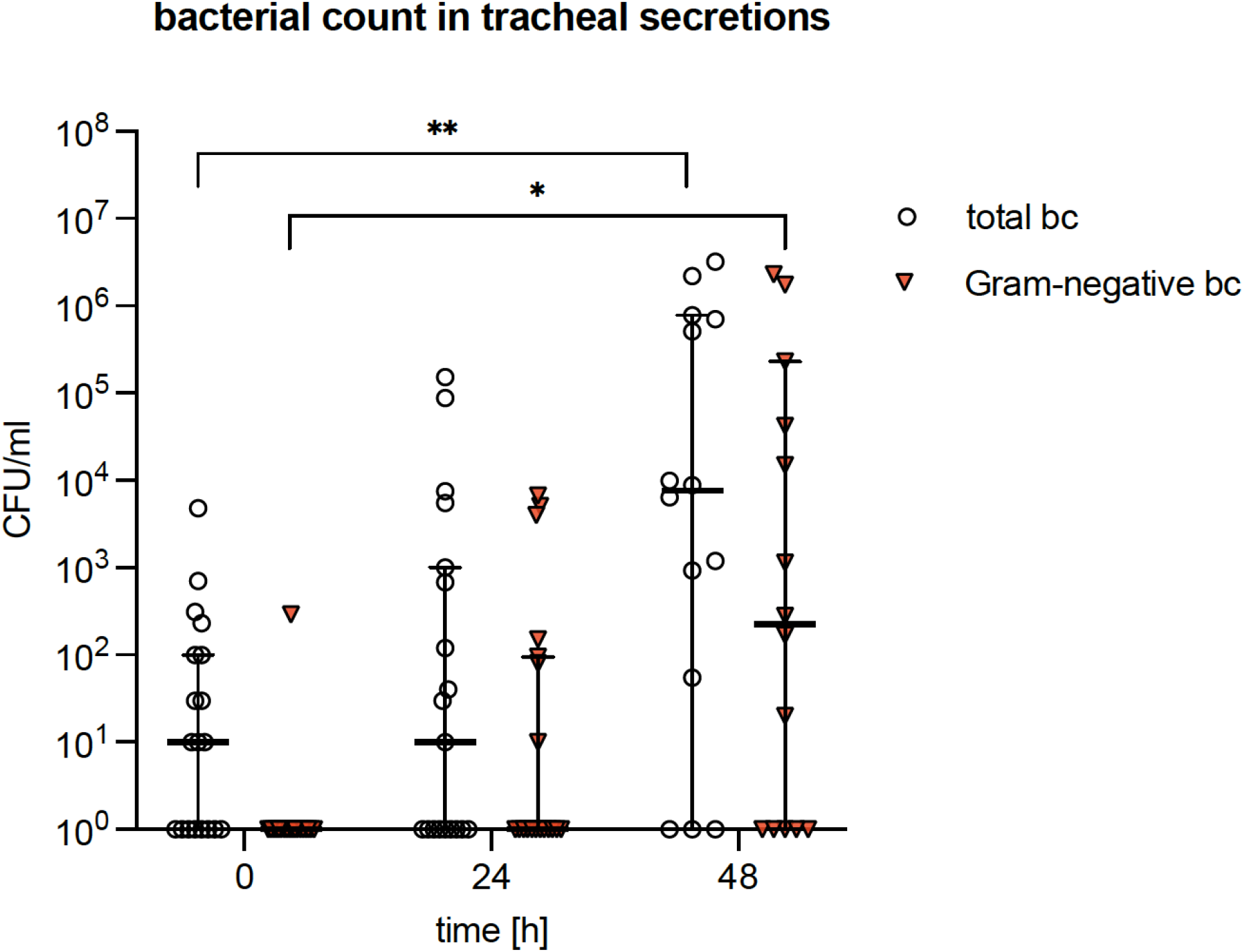
Depicted is the quantitative assessment of total bacterial counts (bc) and Gram-negative organisms of tracheal secretions sampled 1-3 cm below the endotracheal tube. Samples were collected immediately after intubation and after 24 + 1 hours (day1) and 48 + 1 hours (day2) of mechanical ventilation. CFU/ml determinations were carried out through diluting and cultivating respiratory samples on blood agar plates for total bacterial counts and MacConkey agar plates for growth of Gram-negative bacterial species. Shown are median bacterial counts and significant differences (p < 0.05) as determined by Wilcoxon Matched-Pairs Signed-Rank Test.

### Bacterial spread in respiratory secretions during ventilation

To assess the spread of bacteria that occurred within the respiratory tract during ventilation, we analyzed the percentage of tracheal secretions over time without culturable Gram-negative organisms (Tab. 1).

Whereas 94.7% of examined materials were sterile for Gram-negative bacteria at the time of intubation, after 24 hours 36.8% of tracheal secretions showed bacterial growth, e.g. only 63.2% were still sterile. This ratio shifted even further at 48 hours, where in 64.3% of the samples bacterial growth could be detected and only 35.7% remained sterile. To follow the spread of bacteria within the respiratory tract in more detail, we not only identified the detected species, but compared the timepoints of their first occurrences. This analysis revealed that the species we detected in tracheal secretions, were always present in the secretions collected at the cuff. In the vast majority of cases, the detection of specific bacterial species at the cuff occurred 24 to 48 hours prior to their spread to tracheal secretions (Tab. 2).

An effect that even increased over time. Directly after intubation, bacterial colonization of lower tracheal secretions with Gram-negative bacteria affected only one of 19 pigs, where the identical species was found at the cuff. 24 hours later, 85.7% of pigs with positive tracheal secretions grew the same bacterial species, that had already been detected at the cuff at 0h. In the other 14.3% identical species were at least found simultaneously in both secretions. At the 48 hour timepoint, 100% of the cases with colonized tracheal secretions showed the same bacterial species that were found at the cuff at an earlier timepoint. This observation supports the hypothesis that bacterial species found in tracheal secretions during ventilation mainly originate from the microflora at the cuff and spread to lower respiratory areas over time.

### Bacterial species colonizing respiratory secretions during ventilation

To evaluate if bacterial species that are present in respiratory secretions of pigs under ventilation may be relevant for the development of VAP, detailed species determinations by 16S RNA sequencing were carried out. Fig. 1 shows the occurrence rate of species in tracheal secretions of examined pigs. For better clarity the variety of species is categorized in groups. Detailed information can be taken from supplemental Tab.1. The most frequently detected Gram-positive bacterial species were streptococci, often *Streptococcus suis*, that are typical for the porcine physiologic mucosal flora.

For Gram-negative bacteria the most frequently detected species were *Escherichia* spp., occurring in 47.4% of tracheal secretions, followed by other *Enterobacteriaceae* such as *Enterobacter* spp., *Moraxella* spp., *Proteus* spp. and *Klebsiella* spp., *Pseudomonas* spp. and *Acinetobacter* spp., both typical pathogens in ventilator-associated pneumonia, were also frequently detected (Fig. 3). Detailed numbers for species presenting in less than 10% can be found in supplemental Table 1. Overall, we detected the expected residential flora of the upper respiratory tract in addition to many Gram-negative bacterial species that are classical pathogens of VAP.

**Table 1:** Percentage of tracheal secretions without growth of Gram-negative bacterial species on MacConkey agar over 48 hours of ventilation.

|  | time after intubation in hours |  |  |
| --- | --- | --- | --- |
|  | 0 h | 24 h | 48 h |
| Total number of pigs (n) | 19 | 19 | 14 |
| Proportion of sterile tracheal secretions* | 94,7% (18) | 63,2% (12) | 35,7% (5) |
| Proportion of positive tracheal cultures (MacConkey agar)* | 5,3% (1) | 36,8% (7) | 64,3% (9) |

\*Calculated from the total number of animals (n)

**Figure 3:**
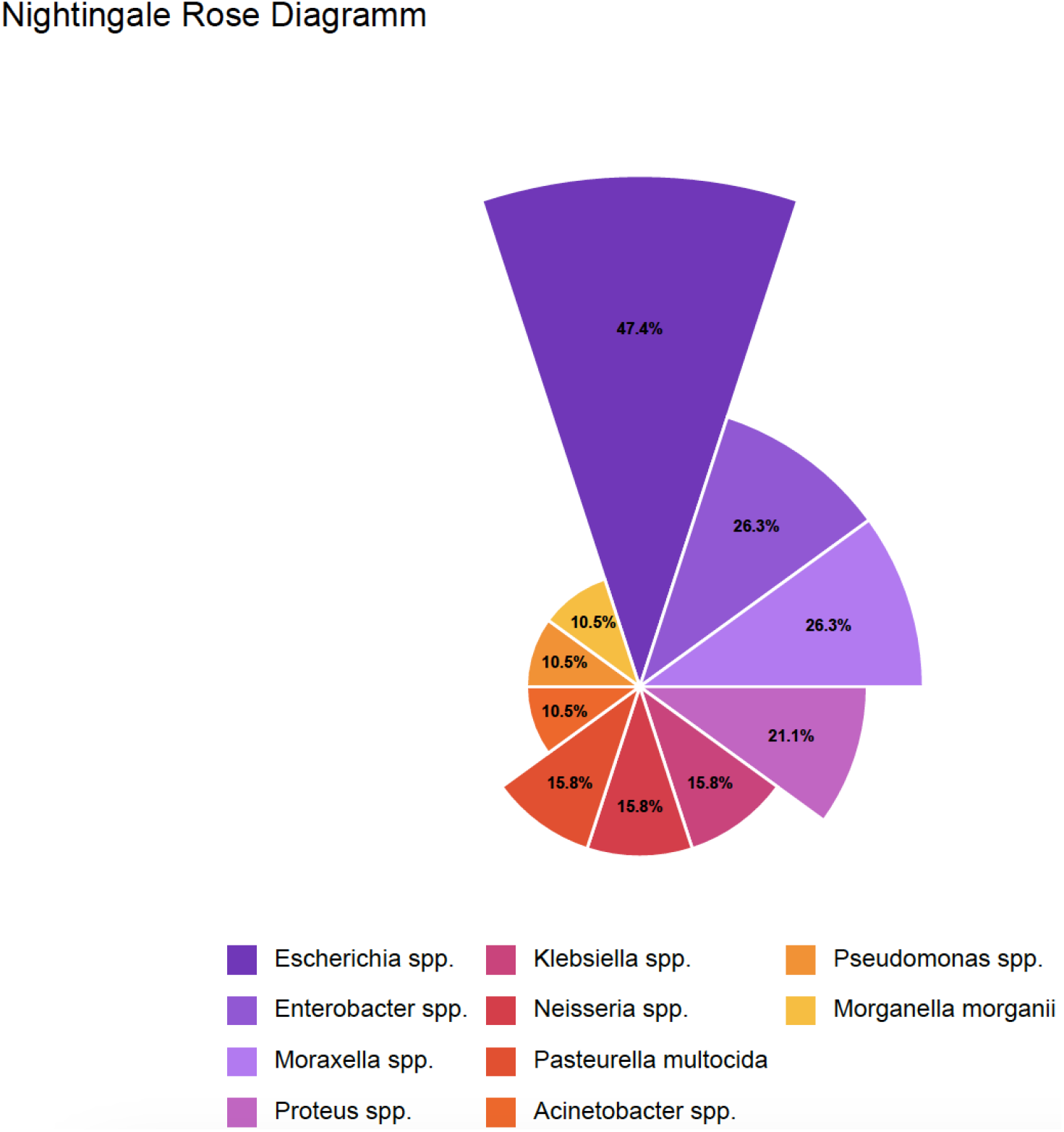
Shown are the frequencies of Gram-negative bacterial species cultivated from tracheal secretions. Following CFU/ml determinations, pure colonies were subcultured and bacterial species determinations were performed by 16S-RNA gene sequencing as detailed in Materials and Methods.

## Discussion

Characterizing the residential flora of respiratory secretions from intubated and ventilated pigs, we noted an extensive increase in bacterial numbers over several orders of magnitude within 24 hours of intubation. This massive increase occurred in particular in the secretions that were collected around the cuff (Fig. 1). Besides the extensive increase in the numbers for all bacterial species, a shift towards and an even higher increase was observed for Gram-negative organisms. Regarding mechanisms that are responsible for the development of VAP the area around the cuff is of particular interest for prophylactic measurements that may lower the incidence of VAP among ventilated patients. At this location natural clearance mechanisms of the trachea are impacted by cuff placement, which interrupts the continuous flow of mucosal secretions towards the oral cavity and thus more or less eliminates the physiological mucociliary clearance (Sackner *et al*., 1975) (Konrad *et al*., 1994). This leads to a pooling and accumulation of pharyngeal secretions above the cuff and creates perfect conditions for bacterial growth. The rise of bacterial numbers by more than three orders of magnitude within 24 hours is exceptional and points to an exponential growth resulting in a bacterial doubling time of about two hours. These conditions resemble bacterial growth how it is typically observed in liquid broth, while it is here obviously occurring *in vivo* in the pooled respiratory secretions at the cuff. Through leakage of these secretions microaspirations can then take place that enable the transition of bacteria from the upper to the lower respiratory tract, where VAP may develop. The importance of cuff leakage for VAP development has repeatedly been discussed (Zolfaghari & Wyncoll, 2011). Focusing local preventive measurements around this area, may thus interfere most effectively with VAP development. The positive effect that subglottical secretion drainage by suctioning the secretions pooled around the cuff has on the incidence of VAP supports this idea (Valles *et al*., 1995) (Dezfulian *et al*., 2005). The hypothesis that the cuff area is crucial in the development of VAP is further supported by following the spread of identical bacterial species within the respiratory tract in our animal model. Comparing the first time of detection of bacterial species that were found in both the cuff secretions and the tracheal secretions, we noted that in almost all cases the specific species was first detected in the cuff area, while in subsequent samples these species were also found in tracheal secretions. This finding appears to indicate an origin of the detected species from the cuff area followed by a spread of these bacteria into the lower tracheal region. Importantly during the 48 hours of ventilation we noticed that 18 of 19 samples were sterile in conventional culture at the time of intubation, whereas after 48 hours of ventilation bacterial growth of Gram-negative bacteria could be detected in 64.3% of these animals (Table 1). Moreover, the Gram-negative bacteria that were found to colonize the tracheal secretions had in almost all cases been detected 24 hours earlier in the secretions obtained from the cuff of the same animal (Table 2). This fact may also offer diagnostic opportunities for the early identification of potential VAP causing bacterial pathogens in clinical settings, if similar observations can be reproduced in studies involving patients.

**Table 2:** Timing of identical Gram-negative bacterial species isolated from cuff and subsequently collected tracheal secretions.

|  | time after intubation in hours |  |  |
| --- | --- | --- | --- |
|  | 0 h | 24 h | 48 h |
| Detection of identical species in tracheal and cuff secretions** | 100% (1) | 100% (7) | 100% (9) |
| Detection of identical species consecutively in tracheal and cuff secretions** | - | 85,7% (6) | 100% (9) |
| Detection of identical species simultaneously in tracheal and cuff secretions** | 100% (1) | 14,3% (1) | 0% (0) |

\*\*Calculated from animals with positive tracheal cultures (MacConkey agar)

Overall, the question remains if the observations of this experimental animal study are of relevance for the development of VAP in humans. Obviously, we did not induce bacterial pneumonia in this model by applying a defined inoculum of a typical nosocomial pathogen to the lower respiratory tract under ventilating conditions. Instead, we observed the shifts in bacterial colonization of respiratory samples occurring under 48 hours of mechanical ventilation. This is clearly a time span that per definition lies prior to the development of VAP, which requires 48 hours of ventilation. So, the actual occurrence of the pneumonia was not in the observed time window, and we cannot be sure if the detection of a bacterial species in the tracheal secretion of pigs would subsequently lead to a VAP. Nevertheless, the pig animal model offers many advantages as pigs are omnivores like humans and their innate and adaptive porcine immune systems more closely resemble that of humans than mice (Meurens *et al*., 2012). Most importantly, the scale and the functional anatomy of the lungs and airways allows clinically relevant models of mechanical ventilation, and intensive-care interventions that will allow to test preventive measurements against VAP in a realistic setting involving endotracheal tubes for example that are also used in humans. Taking into account the bacterial species we found in the respiratory secretions of pigs (Fig. 3), besides classical pig associated bacterial species like *Streptococcus suis* and *Rothia nasimurium*, these included many of the bacterial pathogens, such as ESKAPE pathogens, that are the predominant bacterial species causing human VAP.

Comparing the most common bacterial species in human VAP with the bacterial species we recovered from the respiratory secretions of pigs (Suppl Tab. 1, Fig. 3) we see a striking similarity (Restrepo *et al*., 2013) (Lynch, 2001). The fact that we observed a shift towards these Gram-negative bacterial species in pigs and that they greatly increased in numbers under intubation, which is the key risk factor for VAP development, points in the direction that a similar situation may occur in ventilated human patients. This is supported by previous investigations. While we are not aware of studies quantifying in detail the change of the bacterial load of respiratory secretions occurring before and under ventilation in humans, the shift towards colonizing Gram-negative bacterial species has clearly been demonstrated in the clinical setting (Schwartz *et al*., 1978) (Ewig *et al*., 1999), as well as the importance of respiratory colonization for the overall outcome of patients in the intensive care unit (Dickson *et al*., 2020). We hope therefore, that the current investigation may provide important aspects in understanding the first steps in the development of VAP and that the experimental pig model is an ideal setting to test prophylactic interventions that may limit the extensive increase and spread of VAP associated Gram-negative bacteria in the respiratory tract.

## Acknowledgements

The work of DH, AH, FZ, MH, PR, FM and BS was funded through the BMFTR grant NOVAP, and the DFG (project no. 251 293 561, CRC1149), the work of LS was supported by the Promotionsprogramm Experimentelle Medizin of the Medical Faculty, Ulm University. We thank Stefanie Mauerer and Sandra Kress for expert technical assistance.

## Notes

### Competing Interest Statement

The authors have declared no competing interest.

