## Supplemental Table 1 for "Intubation and mechanical ventilation lead to an extensive expansion of VAP-associated bacterial species in respiratory secretions of ventilated pigs"

Supplemental Table 1: Comprehensive overview of Gram-positive and Gram-negative bacterial species that were cultivated from tracheal secretions

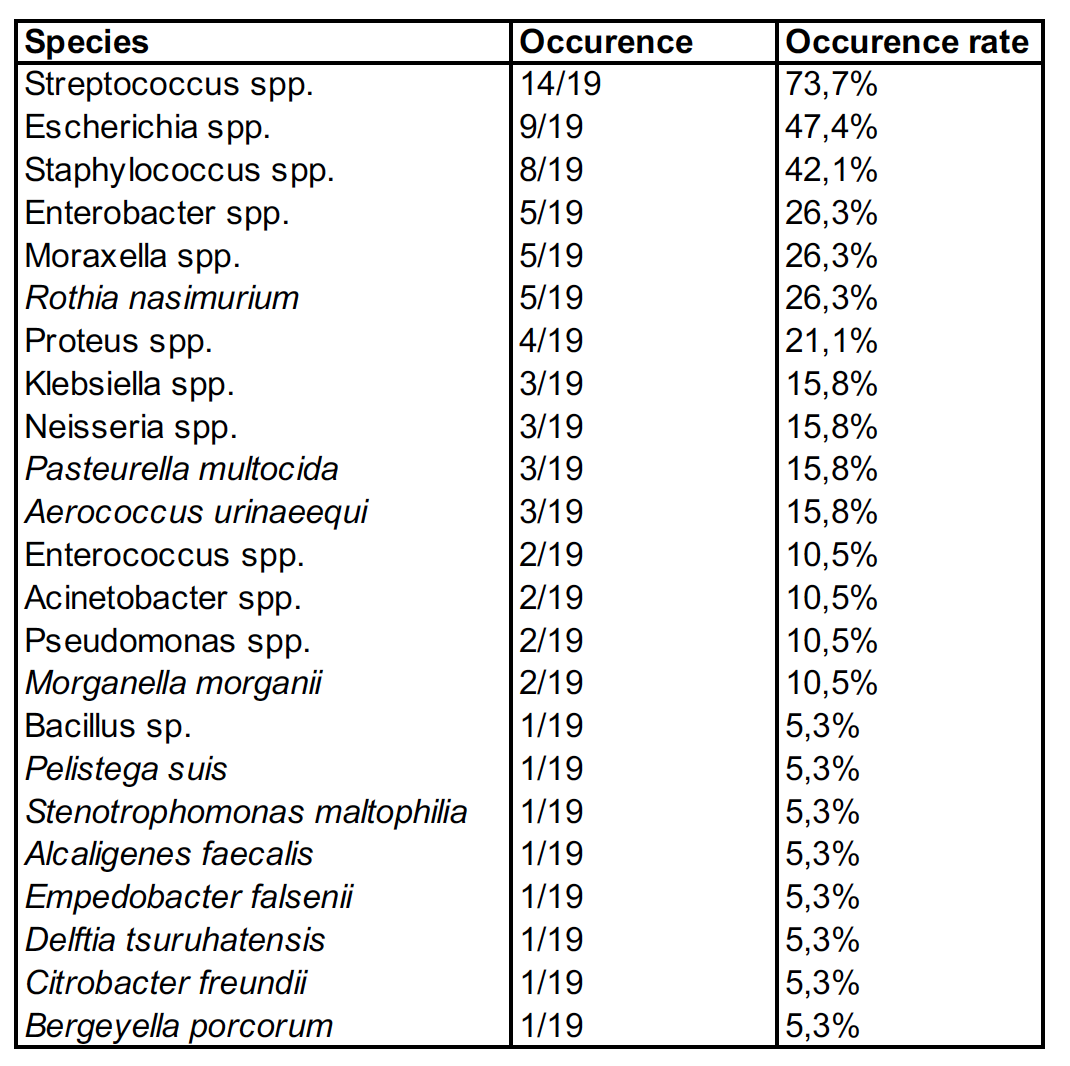
